# *Vibrio cholerae* alkalizes its environment via citrate metabolism to inhibit enteric growth

**DOI:** 10.1101/2022.08.04.502819

**Authors:** Benjamin Kostiuk, Mark E. Becker, Candice N. Churaman, Joshua J. Black, Shelley M. Payne, Stefan Pukatzki, Benjamin J. Koestler

**Affiliations:** Department of Medical Microbiology and Immunology, 6-020 Katz Group Centre, University of Alberta, Edmonton, AB T6G 2S2, Canada; Department of Cell and Developmental Biology, Northwestern University Feinberg School of Medicine, Chicago, IL 60611, USA; Department of Biological Sciences, Western Michigan University, Kalamazoo, Michigan, 49008, USA; Department of Molecular Biology and Genetics, Johns Hopkins University School of Medicine, Baltimore, MD 21205, USA; Department of Molecular Biosciences, The University of Texas at Austin, Austin, Texas, USA; Institute for Cellular and Molecular Biology, The University of Texas at Austin, Austin, Texas, USA; Department of Biology, The City College of New York, New York, NY 10031, USA

**Keywords:** *Vibrio cholerae*, metabolism, citrate, citrate lyase, oxaloacetate decarboxylase, carbonate

## Abstract

*Vibrio cholerae* is a Gram-negative pathogen, living in constant competition with other bacteria in both marine environments and during human infection. One competitive advantage of *V. cholerae* is the ability to metabolize diverse carbon sources such as chitin and citrate. We observed that when *V. cholerae* strains were grown on a medium with citrate, the medium’s chemical composition turned into a hostile alkaline environment for Gram-negative bacteria such as *Escherichia coli* and *Shigella flexneri*. We found that although the ability to exclude competing bacteria was not contingent on exogenous citrate, *V. cholerae* citrate metabolism mutants Δ*oadA*-1, Δ*citE*, and Δ*citF* mutants were not able to inhibit *S. flexneri* or *E. coli* growth. Lastly, we demonstrated that while the *V. cholerae* mediated increased medium pH was necessary for the enteric exclusion phenotype, secondary metabolites such as bicarbonate (protonated to carbonate in the raised pH) from the metabolism of citrate enhanced the ability to inhibit the growth of *E. coli*. These data provide a novel example of how *V. cholerae* outcompetes other Gram-negative bacteria.

## Introduction

For bacteria to thrive in a competitive environment, they must be highly effective in resource acquisition to proliferate their niche^1,2^. Bacteria employ both passive and active forms of competition^2–4^. Active processes include the secretion of toxins or the sequestration of resources^1,3^. Passive mechanisms include the secretion of waste products of their secondary metabolic pathways, making the environment hostile for their competitors; such secondary metabolites are not required for the producing organism and are referred to as allelochemicals^5^.

*Vibrio cholerae*, the causative agent of the diarrheal disease cholera, is a Gram-negative pathogen that primarily resides in marine reservoirs, and causes disease upon human ingestion^6^. Because the pathogenic cycle of *V. cholerae* involves transitioning between its natural marine environment and the human host, *V. cholerae* has evolved to be highly competitive in both of these environments^6^. During these transitions, *V. cholerae* interacts with many different types of microbial organisms, including various eukaryotes and bacteria of the same or other species^7,8^. As a consequence of residing in diverse microbial communities, *V. cholerae* has evolved multiple competitive mechanisms that are effective against other members of its species, other bacterial species, or eukaryotic predators^1,3,7^.

One such *V. cholerae* survival mechanism is using multiple carbon sources for energy^9^. A prominent example is *V. cholerae’s* ability to use chitin as a carbon source, as chitin is abundant in marine environments^10,11^. In addition to chitin, *V. cholerae* can metabolize other carbon sources, such as dietary citrate, that some of its competitors cannot utilize^12^. In addition, strains of *V. cholerae* that successfully colonize humans endure the low pH stress of the stomach and small intestine^9,13^. *V. cholerae’s* ability to thrive in multiple environments has given rise to novel mechanisms of competition. For example, *V. cholerae* actively secretes vibriobactin, and this siderophore sequesters iron to provide this essential nutrient for itself and prevent other species from using it^1^.

*V. cholerae* is responsible for seven recorded pandemics. Pandemic strains are divided into two biotypes; the seventh pandemic *V. cholerae* O1 El Tor biotype, and the sixth pandemic O1 classical biotype, which evolved as distinct lineages^14–18^. Differences in *V. cholerae* metabolism profiles are implicated in significant differences in intraspecies fitness. Notably, some El Tor strains produce 2,3-butanediol when metabolizing glucose; this creates a more favorable environment for survival of El Tor strains than classical *V. cholerae* pandemic strains, which generate an unsuitably low pH when grown in the presence of glucose^18,19^. Here, we investigated whether *V. cholerae* uses its ability to metabolize citrate for growth advantages, and present a model for a competition mechanism in which *V. cholerae* defines its chemical microenvironment through the metabolism of citrate. Using a simple in-vitro assay, we show that *V. cholerae* metabolites create an environment hostile to competing bacteria through the increase in pH and bicarbonate production.

## Results

### V. cholerae inhibits the growth of enteric bacteria

The *V. cholerae* El Tor strain C6706, which is contributing to the ongoing 7^th^ pandemic^20^, suppresses the growth of other microorganisms such as *Escherichia coli* in a cross-streaking assay on citrate-containing media when grown for an extended time (54 hours)^21^. A cross-streaking assay involves growing a streak of *V. cholerae* down the center of an agar plate. Following bacterial growth, bacteria are scraped off. The plate is subjected to chloroform vapor to remove residual bacteria, and a single line of indicator strain like *E. coli* is streaked perpendicular to the original bacterial growth. The plates are incubated overnight, and the growth of the indicator strain is recorded. A streak of *V. cholerae* C6706 grown on Nutrient Broth agar supplemented with citrate (C.B.), but not on Nutrient Broth agar, resulted in inhibition of the growth of *E. coli* or *Shigella flexneri*^22^. Because *V. cholerae* is no longer present on the agar plate when *E. coli* inhibition occurs, the interpretation was that growth on citrate stimulated *V. cholerae* secretion of an unknown factor.

Two groups had previously proposed that this is a competitive behavior where an unidentified bacteriocin-like compound with bactericidal activity against *E. coli* is secreted^21,23^. A third group suggested a metabolic byproduct was responsible for this growth inhibition^22^. We confirmed this phenotype and found that prolonged growth (48 hours) of *V. cholerae* C6706 on L.B. agar before cross-streaking also resulted in growth inhibition of the closely related enteric bacteria *S. flexneri*, suggesting that the effect was stimulated by, but not dependent on, citrate present in the media (Fig. 1A). We also performed a supernatant inhibition assay, where *V. cholerae* C6706 supernatants were centrifuged and filter-sterilized to remove bacteria and debris, and then we measured the ability of the supernatant to prevent *E. coli* or *S. flexneri* growth; this allowed us to quantitatively compare the inhibition properties of *V. cholerae* C6706 by calculating the half-maximal inhibitory concentration (IC_50_). We found that after growth in L.B. broth, *V. cholerae* C6706 cell-free supernatants had dose-dependent inhibitory activity against *S. flexneri*, whereas *S. flexneri* cell-free supernatants did not impede the growth of *S. flexneri* (Fig. 1B).

**Figure 1.**
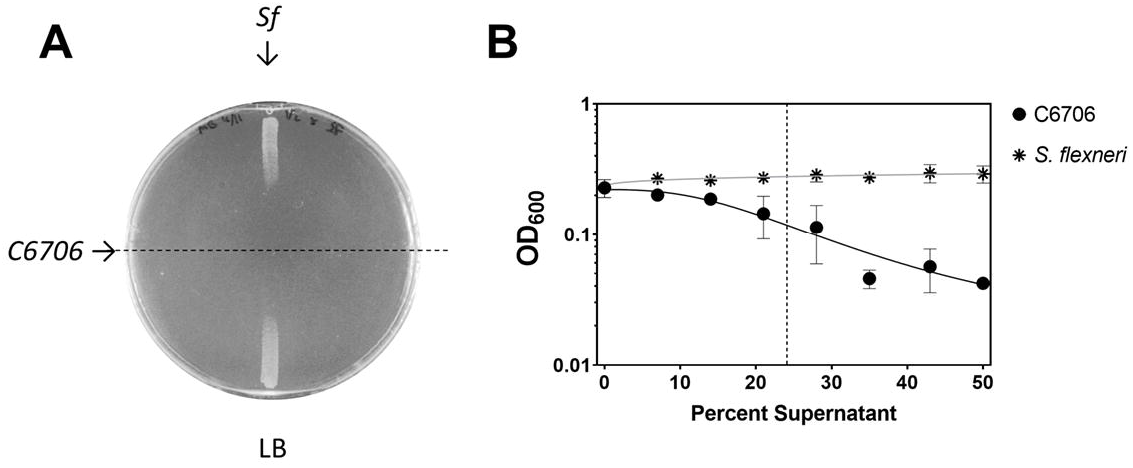
Extended culture of *V. cholerae* C6706 inhibits enteric growth. (A) A cross-streak assay demonstrates that *V. cholerae* C6706 inhibits the growth of *S. flexneri* CFS100. *V. cholerae* C6706 was grown for 72 hours on L.B. agar. Bacteria were scraped off the plate, followed by chloroform vapor treatment to kill residual cells. *S. flexneri* CFS100 was then streaked perpendicular to the *V. cholerae* growth, and incubated for approximately 12 hours. There was a clear zone of inhibition proximal to where the *V. cholerae* was originally located. (B) The supernatant inhibition assay demonstrates that *V. cholerae* cell-free supernatants inhibit *S. flexneri* CFS100 growth. *V. cholerae* C6706 or *S. flexneri* CFS100 was grown in L.B. broth for 72 hours. Cultures were then centrifuged, supernatants were collected, and filtered through a 0.022 µm PVDF filter. Supernatants were then diluted in L.B. at different ratios. The ability of *S. flexneri* CFS100 to grow in these supernatants was determined by measuring the OD_600_ after 6 hours. *V. cholerae* supernatants inhibited *S. flexneri* growth in a dose-dependent manner, whereas *S. flexneri* supernatants had no inhibitory effects on *S. flexneri* growth. The IC_50_ (indicated by the dotted line) of *V. cholerae* supernatants was calculated to be 24.1%. Each point shows the mean of 3 replicates, error bars show standard deviation. Trendlines show a nonlinear regression (log inhibitor vs response, 4 parameter).

We sought to determine if the inhibitory effect of *V. cholerae* C6706 supernatants on *S. flexneri* growth in liquid media were bacteriostatic or bactericidal in nature. We grew *S. flexneri* in various concentrations of supernatant derived from the *V. cholerae* C6706 strain and *S. flexneri*, and measured growth (OD_600_) over time. We then determined the CFS100 growth rate and lag times in each of these conditions. We observed a modest effect on the growth rate of *S. flexneri* when grown in various concentrations of supernatant from either strain (Fig. 2A); however, there was a significant dose-dependent increase in the lag time prior to *S. flexneri* growth in *V. cholerae* C6706 supernatant, relative to *S. flexneri* grown in equivalent concentrations of *S. flexneri* supernatant (Fig. 2B), suggesting that this inhibitory effect is largely bacteriostatic in nature.

**Figure 2.**
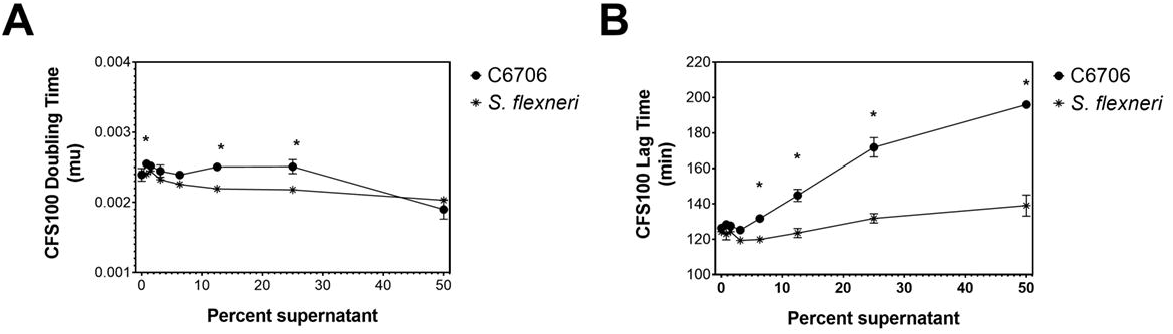
Effect of *V. cholerae* C6706 supernatants on *S. flexneri* growth. The supernatant inhibition assay was performed using supernatants derived from *V. cholerae* C6706 or *S. flexneri* CFS100, and the growth of *S. flexneri* CFS100 was monitored by OD_600_ measurements. The growth rate and lag time of *S. flexneri* CFS100 grown in different concentrations of supernatant derived from *V. cholerae* C6706 or *S. flexneri* CFS100 was quantified using the R grofit package^55^. Each growth curve was replicated in triplicate. (A) We observed a modest increase in doubling time caused by *V. cholerae* C6706 supernatant, relative to supernatant derived from *S. flexneri* CFS100. (B) We observed a significant, dose-dependent increase in lag time in *S. flexneri* CFS100 growth caused by *V. cholerae* C6706 supernatants, relative to the *S. flexneri* CFS100 supernatants.

### Citrate metabolism contributes to V. cholerae inhibition of enteric bacteria

To determine if the citrate-stimulated competition mechanism of the El Tor strain C6706 is a common trait of *V. cholerae*, we investigated whether other strains share this phenotype. We first determined the ability of 15 *V. cholerae* strains to grow on citrate using Simmons’ citrate agar^24^; this test relies on an organism to grow using citrate as a sole carbon source. The strains used in this study included both environmental and pandemic-causing strains^25^. Surprisingly, we found heterogeneity in not only the ability of various *V. cholerae* strains to create a hostile environment, but also on the ability of *V. cholerae* strains to grow using citrate. While all strains were able to grow in L.B. or C.B., we found that 9 of the 15 *V. cholerae* strains could grow on Simmons’ citrate agar (Table 1; green row). Of those 9, only five were able to create a hostile environment for *E. coli* in a cross-streak assay (Table 1; blue row). While *E. coli* cannot grow on this conditioned media, *V. cholerae* C6706 can grow in the conditioned media. Some *V. cholerae* strains (listed in the red row) grew on citrate provided as the sole carbon source yet were not able to inhibit the growth of *E. coli*. We did not observe any notable correlation between the source of the *V. cholerae* strains (i.e pandemic or environmental) and the ability to create a hostile growth environment for *E. coli*^25^.

**Table 1.** *V. cholerae* strains differ in their ability to metabolize citrate and prevent *E. coli* growth in a cross-streaking assay. Fifteen *V. cholerae* strains analyzed in this study can be grouped into three distinct groups based on their ability to metabolize citrate as well as their ability to prevent the growth of *E. coli* in a cross-streaking assay on C.B. The blue group contains those strains both able to grow on Simmons’ citrate agar and prevent *E. coli* growth. The red group contains those strains able to grow on Simmons’ citrate agar but not prevent the growth of *E. coli*. Finally, the green group contains strains not able to grow on Simmons’ citrate agar, and subsequently not able to prevent the growth of *E. coli*. Analysis was replicated three times.

| *V. cholerae*<br>producing strain | Growth on<br>Simmons' agar | pH of liquid LB<br>medium – 48 hours | Cross-streak assay |  |
| --- | --- | --- | --- | --- |
|  |  |  | Growth of<br>*E. coli* MG1655 | Growth of<br>*V. cholerae* C6706 |
| C6706 | + | 8.6 ± 0.9 | - | + |
| C6706 $\Delta citE::Tn$ | + | n.d. | + | + |
| C6706 $\Delta citF::Tn$ | + | 8.2 ± 1.1 | + | + |
| N16961 | + | 7.5 ± 1.2 | - | + |
| O395 | + | 6.7 ± 0.1 | - | + |
| DL4211 | + | 9.1 ± 0.2 | - | + |
| 1587 | + | 7.6 ± 1.2 | - | + |
| V52 | + | 7.5 ± 1.2 | + | + |
| MZO-3 | + | 6.7 ± 0.0 | + | + |
| 27-4080 | + | 7.6 ± 1.0 | + | + |
| MAK-757 | + | 7.2 ± 0.1 | + | + |
| AM19226 | - | 7.3 ± 0.0 | + | + |
| V51 | - | 7.6 ± 1.0 | + | + |
| NIH41 | - | 6.9 ± 0.0 | + | + |
| MZ02 | - | 7.1 ± 0.0 | + | + |
| C6709 | - | 7.6 ± 1.0 | + | + |
| CA401 | - | 7.0 ± 0.0 | + | + |

Very few studies have experimentally examined the basis of *V. cholerae* citrate metabolism^26^. *V. cholerae* C6706 encodes homologous genes associated with the citric acid cycle (TCA) and also citrate fermentation; in this pathway, a sodium/citrate symporter facilitates citrate uptake, and then a citrate lyase (ACLY) cleaves citrate into acetate and oxaloacetate, the latter of which is converted to CO_2_ and pyruvate by oxaloacetate decarboxylase^27–30^ (Fig. 3). To determine if citrate fermentation contributes to enteric growth inhibition, we tested two *V. cholerae* C6706 transposon mutants with disrupted genes encoding ACLY, Δ*citE*::Tn and Δ*citF*::Tn, to determine the importance of citrate metabolism pathway on the *V. cholerae* C6706 competition phenotype^31^. Although both these mutants still displayed growth on Simmons’ citrate agar, they did not prevent *E. coli* growth (Table 1). Similarly, a mutation in *oadA-1*, coding for oxaloacetate decarboxylase, which blocks the conversion of oxaloacetate to pyruvate in citrate metabolism^28^, also eliminated the inhibitory activity produced by *V. cholerae* C6706 in a supernatant inhibition assay (Fig. 4A). We concluded that the inhibitory activity of *V. cholerae* C6706 on *S. flexneri* and *E. coli* is dependent on *V. cholerae* C6706 citrate fermentation.

**Figure 3.**
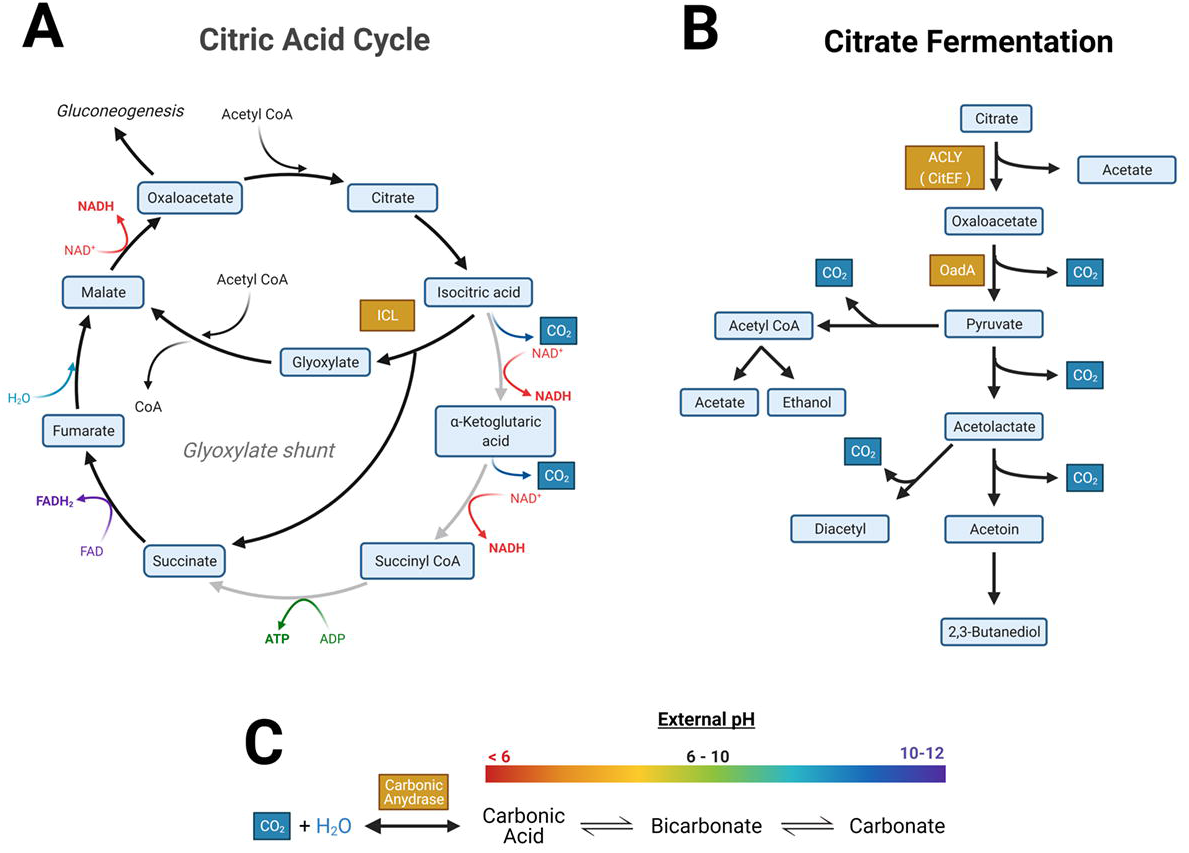
Conceptual model summarizing that the metabolism of citrate under basic conditions favors the formation of carbonate. (A) Diagram of the Citric Acid Cycle in *V. cholerae*, highlighting the glyoxylate shunt. (B) Abridged metabolic flow diagram showing the fermentation of citrate to 2,3-Butanediol or Acetyl-coA, producing CO_2_ as a byproduct (in blue). (C) CO_2_ is converted to carbonic acid through the enzyme carbonic anhydrase. An equilibrium exists between carbonic acid, bicarbonate and carbonate. More basic conditions favor the production of the negatively charged carbonate. Enzymes of interest in this study are highlighted in orange. Other enzymes and other cofactors have been omitted to emphasize the production of CO_2_. Created with BioRender.com.

**Figure 4.**
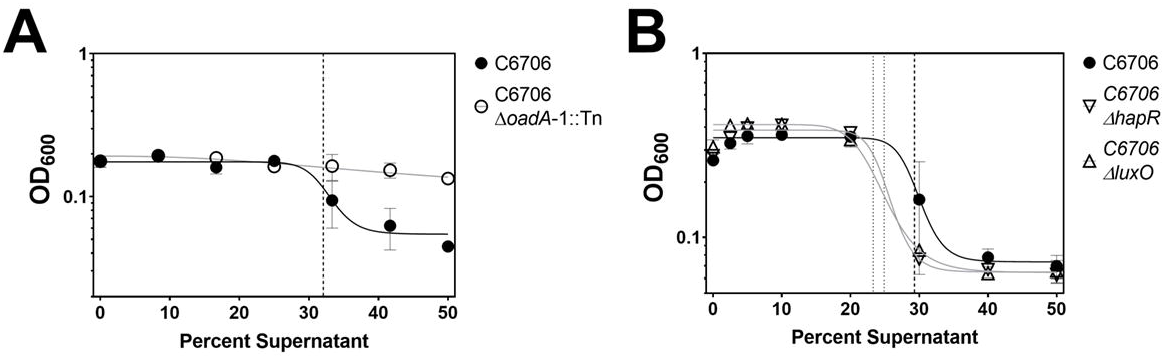
Quantification of the *V. cholerae* supernatant inhibitory activity of *S. flexneri* using a cell-free assay. (A) Disruption of a *V. cholerae* oxaloacetate decarboxylase (*oadA-1*) eliminates supernatant inhibitory activity. Supernatants derived from the growth of the *V. cholerae* Δ*oadA*-1::Tn mutant were unable to inhibit *S. flexneri* CFS100 growth, compared to the W.T. strain. The IC_50_ of the W.T. strain is shown by the dashed line. (B) Disruption of *V. cholerae* quorum sensing master regulators does not significantly alter supernatant inhibition activity. Supernatants derived from the growth of *V. cholerae* Δ*hapR*::Tn or Δ*luxO*::Tn mutants were able to inhibit the growth of *S. flexneri* CFS100 similar to the *V. cholerae* C6706 WT strain. Each point is the mean of 3 replicates, and error bars show standard deviation. Trendlines show a nonlinear regression (log inhibitor vs response, 4 parameter). IC_50_ values are shown by the dashed (W.T.) and dotted (Δ*hapR* & Δ*luxO*) lines.

### V. cholerae raises pH of media during growth

We sought to characterize the nature of the *V. cholerae* C6706 inhibition of *S. flexneri*. The secretion of inhibitory factors is a bacterial behavior that often conveys a communal fitness advantage; therefore, we hypothesized that the *V. cholerae* inhibitory mechanism was regulated by quorum sensing. However, we found that there was no significant difference in the IC_50_ of W.T. *V. cholerae* C6706 supernatants (29.3, dashed line) compared to Δ*hapR* (24.9 dotted line) and Δ*luxO* (23.3, dotted line) quorum-sensing mutant strains in our supernatant inhibition assay (Fig. 4B). We also investigated the hypothesis that the *V. cholerae* C6706 secreted factor is a protein^21,23^. If the *V. cholerae* C6706 secreted factor were a protein, it might be sensitive to heat; however, *V. cholerae* C6706 cell-free supernatants that were incubated at 60 °C for 60 minutes still retained their ability to inhibit *S. flexneri* growth (Table 2). Furthermore, proteinase K treatment of *V. cholerae* C6706 cell-free supernatants also was not able to significantly alleviate *S. flexneri* growth inhibition, as well as filtration through a 1 kDa filter (Table 2). These data together do not support the hypothesis that the *V. cholerae* C6706 secreted inhibitory factor is a protein.

**Table 2.**
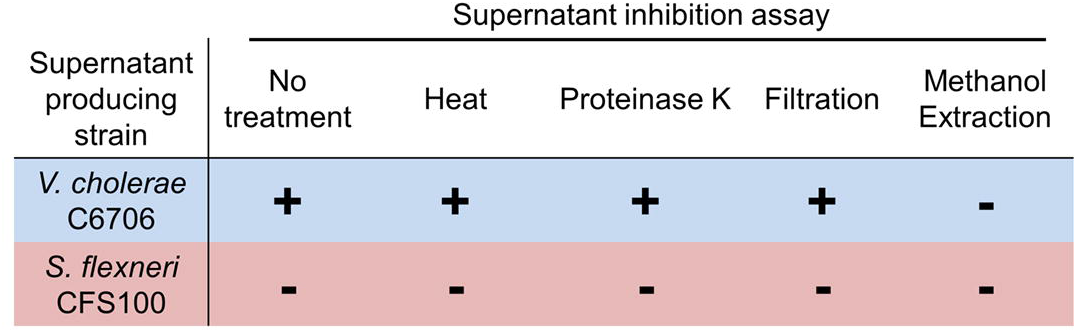
Supernatant treatment effects on the ability to inhibit *S. flexneri* CFS100 growth. *V. cholerae* C6706 or *S. flexneri* CFS100 were grown for 72 hours in L.B. broth, cells were removed, and then cell-free supernatants were treated and used in a supernatant inhibition assay to inhibit the growth of *S. flexneri* C6706. Treatments included heat treatment (60° C for 1 hour), treatment with proteinase K, filtration through a 1-kDa filter, or methanol phase separation followed by reconstitution in saline. + indicates supernatants were able to inhibit the growth of *S. flexneri* CFS100, whereas – indicates that supernatants were not able to inhibit the growth of *S. flexneri* CFS100.

Supernatant inhibition assay
| Supernatant producing strain | No treatment | Heat | Proteinase K | Filtration | Methanol Extraction |
| --- | --- | --- | --- | --- | --- |
| *V. cholerae* C6706 | + | + | + | + | - |
| *S. flexneri* CFS100 | - | - | - | - | - |

We next investigated the hypothesis that the *V. cholerae* C6706 secreted inhibitory factor is a metabolic byproduct^22^. We performed a methanol extraction to isolate metabolites from *V. cholerae* C6706 cell-free supernatants. Methanol was added to *V. cholerae* C6706 cell-free supernatants at a ratio of 6:1 to precipitate protein. The liquid fraction was collected and evaporated by vacuum centrifugation; the remaining content was resuspended in saline. We found that metabolites extracted from *V. cholerae* C6706 cell-free supernatants did not display any inhibitory activity towards *S. flexneri* (Table 2). As part of this experiment, we measured the pH of the supernatants before and after extraction, and the pH of metabolites resuspended in saline after methanol extraction. Prior to culture, the pH of the L.B. broth used to grow *V. cholerae* C6706 and *S. flexneri* was approximately 7.0; however, the pH of *V. cholerae* C6706 cell-free supernatants after 48-hours of culture was approximately 9.5, consistent with prior studies^22^. After methanol extraction, the pH of *V. cholerae* C6706 cell-free supernatant metabolites resuspended in saline was approximately 7.0. We also measured the pH of C.B. broth before and after *V. cholerae* C6706 growth. Like L.B. broth, the pH of liquid C.B. was approximately 7.0 before inoculation; following 48-hour growth of C6706 in C.B., the pH rose by approximately two logs to 9.0. To determine if other *V. cholerae* strains raise media pH similar to the C6706 strain, we grew each of our *V. cholerae* strains in L.B. for 48 hours, and then quantified the pH of cell-free supernatants. Only the *V. cholerae* C6706 and DL4211 strains demonstrated significantly elevated supernatant pH (Table 1).

### V. cholerae C6706 alkalization of media inhibits S. flexneri growth

A previous study reported that *E. coli* and *S. flexneri* do not grow when the external pH is higher than 9.0^32^. To confirm that *S. flexneri* is also sensitive to basic conditions, we grew *S. flexneri* in L.B. adjusted to different pH. We observed a modest growth defect when the pH of L.B. was brought to 9.0, and we observed no growth when the pH was raised to 9.5 and 10.0 (Fig. 5A). Because the pH of *V. cholerae* C6706 supernatants was higher than 9.0 after 72 hours, we hypothesized that alkaline pH contributed to the inhibition of *S. flexneri* growth; we tested this hypothesis by adjusting the pH of *V. cholerae* C6706 cell-free supernatants. When the pH of *V. cholerae* C6706 cell-free supernatants was adjusted to 7.5 using HCl, we found that *V. cholerae* C6706 cell-free supernatants could not inhibit *S. flexneri* growth in a liquid inhibition assay (Fig. 5B).

**Figure 5.**
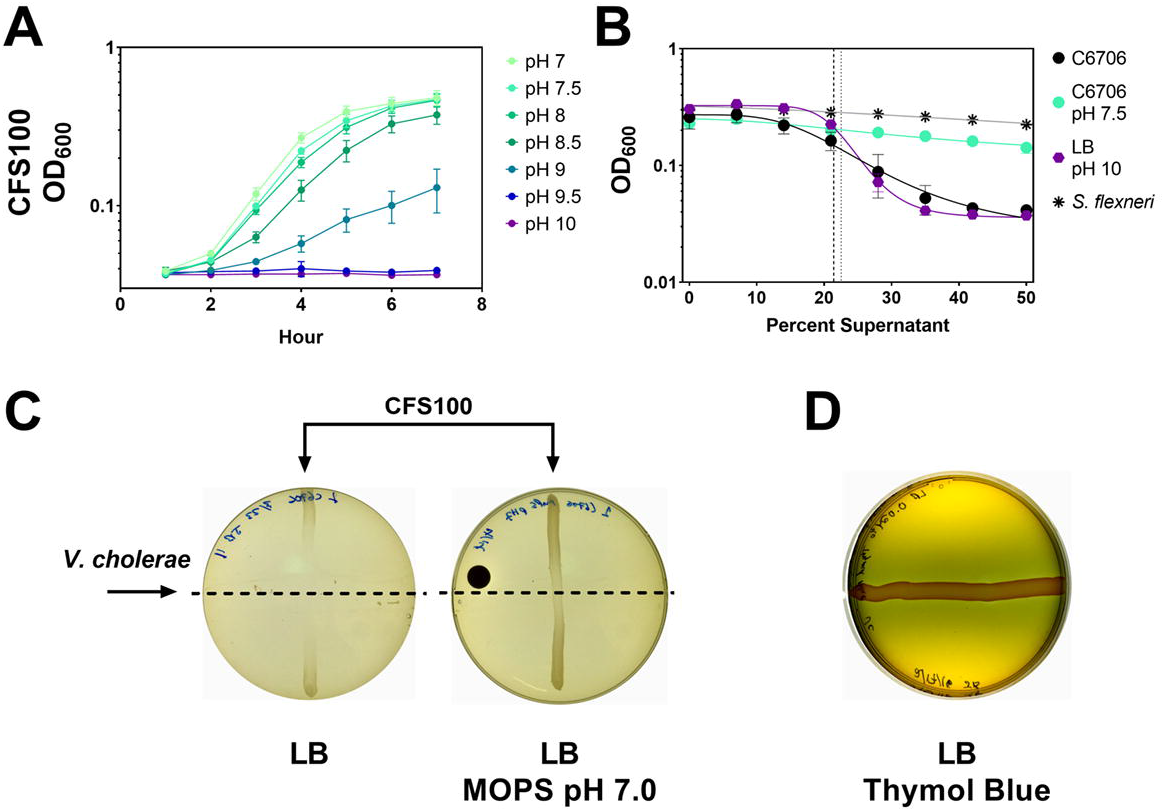
Media alkalization is necessary for *V. cholerae* supernatant inhibitory activity. (A) Alkaline pH inhibits *S. flexneri* CFS100 growth. L.B. was adjusted to different pH using sodium hydroxide, and *S. flexneri* growth was quantified over time by measuring OD_600_. When the media pH exceeded 9.0, significant *S. flexneri* growth inhibition occurred. (B) Changes in pH correspond to *V. cholerae* supernatant inhibitory activity. The supernatant inhibition assay was used to quantify the inhibitory activity of *V. cholerae* supernatants with adjusted pH. When the pH of *V. cholerae* was lowered to 7.5, no *S. flexneri* growth inhibition was observed; in contrast, when the pH of L.B. was raised to 10.0, the pattern of growth inhibition was similar to that of *V. cholerae* supernatants. (C) Buffering L.B. agar plates reduces *V. cholerae* supernatant inhibitory activity. L.B. agar was prepared with and without MOPS buffer (pH 7.0, 50 mM). When the cross-streak assay was performed, there was minimal zone of inhibition observed in the MOPS plate, compared to the L.B. agar plate. (D) Thymol blue was supplemented to L.B. agar to visualize pH changes after V. cholerae growth. After 24 hours, a blue coloration change is observed surrounding the V. cholerae streak corresponding with the zone of inhibition we observe, indicating an increase in media pH.

Furthermore, when the pH of L.B. was raised to 10.0, it had the same inhibitory effect towards *S. flexneri* as *V. cholerae* C6706 supernatants (Fig. 5B). To determine if this same effect occurred in our cross-streaking assay, we grew *V. cholerae* C6706 on L.B. agar buffered with MOPS (50 mM) at pH 7.0. When we performed our cross-streaking assay using this buffered medium, we observed little to no inhibition of *S. flexneri* growth (Fig. 5C). We also visualized *V. cholerae* C6706 alkalization of L.B. agar by adding Thymol Blue, a pH indicator that transitions from yellow to blue between pH 8.0 and 9.2. After 24 hours, we observed a blue zone surrounding *V. cholerae* C6706 growth, which corresponded to the *S. flexneri* zone of inhibition we observed (Fig. 5D).

Similar to growth in L.B., when *V. cholerae* C6706 was grown in C.B., we found that *V. cholerae* C6706 cell-free supernatants did not inhibit *E. coli* growth after pH was re-adjusted to ∼7.0 with HCl, or if the *V. cholerae* C6706 growth medium was buffered to prevent the increase in pH (Fig. 6A). Carbonate is a byproduct of citrate metabolism under basic conditions and is known to prevent the growth of *E. coli*^33^, thus we hypothesized that *V. cholerae* C6706 metabolism of citrate generates carbonates and basic conditions responsible for deprotonating bicarbonate to carbonate (Fig. 3). To test the hypothesis, we examined *E. coli* and *V. cholerae* C6706 growth in a combination of sodium bicarbonate and a basic pH, mimicking the conditions we anticipate are caused by *V. cholerae* C6706 grown in citrate-containing media. We found that these conditions restricted the growth of *E. coli* more profoundly than basic pH or bicarbonate alone (Fig. 6B). *V. cholerae* C6706 grew under all conditions, and even though *V. cholerae* C6706 grew more slowly in the combination of high pH and bicarbonate (Fig. 6B), it still reached the mid-logarithmic phase of growth. The supplementation of bicarbonate to media at a pH of 9.0 reduced the growth rate of *E. coli*, but not *V. cholerae* (Fig. 6C).

**Figure 6.**
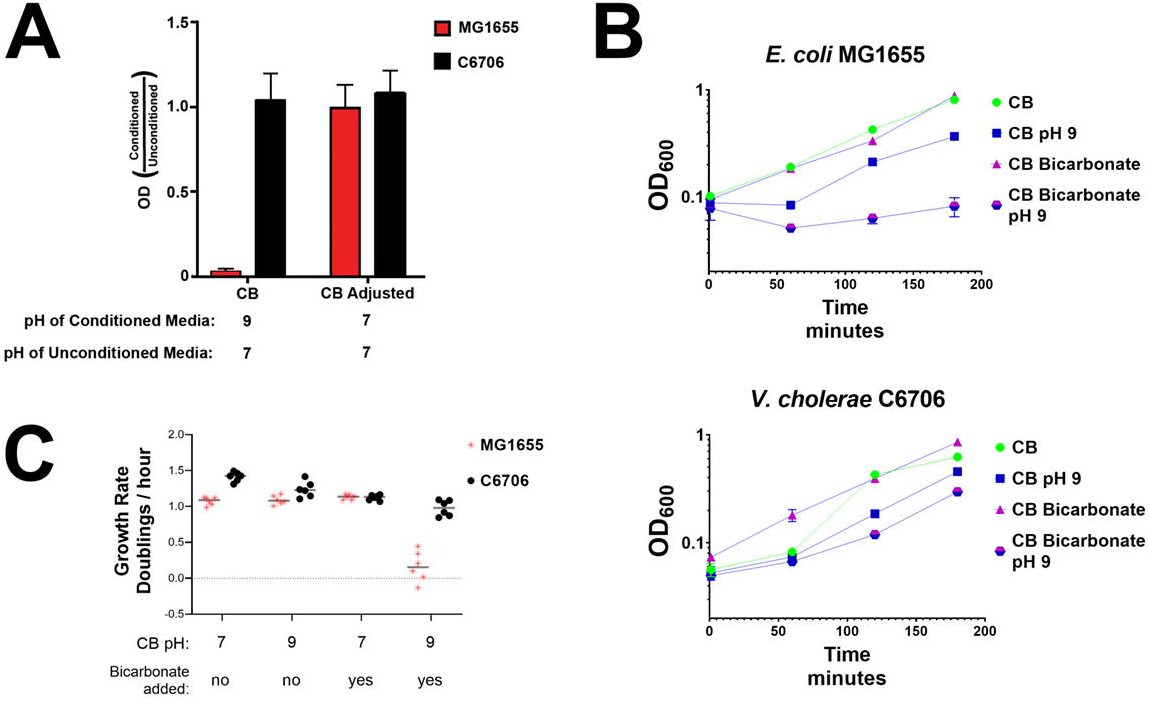
The growth inhibition of *E. coli* can be replicated in a *Vibrio-*free system using a high pH and bicarbonate. (A) Conditioned C.B. media by growth of *V. cholerae* prevents the growth of *E. coli* in a pH-dependent manner. Optical density of indicated bacterial culture after four hours of growth in different conditioned media compared to unconditioned media. *V. cholerae* C6706 was grown in C.B. media for 48 hours. The pH cell-free supernatant was then either adjusted to 7.0 or left as indicated. The relative ability of both *E. coli* and *V. cholerae* to grow in this conditioned media was determined by the optical density after four hours compared to the optical density in unconditioned media. (B) Preventing the growth of *E. coli* can be mimicked with pH and bicarbonate. C.B. media is either supplemented with bicarbonate, or not and either left at pH 7 or artificially raised to a pH of 9. Top shows the growth of *E. coli* MG1655 in different media conditions, bottom shows the growth of *V. cholerae* C6706. (C) Growth rates of both *E. coli* and *V. cholerae* were determined in these four media by taking the slope of the linear portion of the growth curve. The y-axis represents the growth rate of *E. coli* divided by the growth rate of *V. cholerae* under the conditions indicated on the x-axis.

## Discussion

It was first reported some fifty years ago that *V. cholerae* strains could inhibit the growth of enteric bacteria, but the mechanism of this inhibition was never determined^21–23^. While more recent studies have illustrated how *V. cholerae* kills other bacteria using a Type VI secretion system (T6SS)^3^, the mechanism first described by Chakrabarty et al. is a contact-independent mechanism^21^. Here, we investigated the hypotheses originally posited by these groups regarding the *V. cholerae* mechanism of contact-independent inhibition of enteric growth^21–23^. Our findings were consistent with the findings of Bhaskaran et al., who proposed that secreted carbonates raised media pH and caused enteric growth inhibition^22^. We provide additional support of this hypothesis by demonstrating that *V. cholerae* C6706 mutants defective in the conversion of citrate to oxaloacetate (*citE, citF*) and the conversion of oxaloacetate to pyruvate (*oadA-1*) were not able to inhibit enteric growth (Table 1, Fig. 4A). We cannot rule out the possibility that *V. cholerae* C6706 produces a bacteriocin-like protein to inhibit the growth of enteric bacteria; however, our data suggest that such a protein would require an alkaline environment to function, as buffering media pH or re-adjusting *V. cholerae* C6706 cell-free supernatants to a neutral pH abolishes inhibitory activity (Fig. 5 & 6).

Carbohydrate metabolism is an essential aspect of *V. cholerae* pathogenesis. Previous studies have shown that *V. cholerae* relies on standard carbon metabolism pathways coupled with oxygen respiration in the host, including the Embden-Meyerhof-Parnas (glycolysis) and Entner-Doudoroff pathways^34–36^. Recent studies demonstrate that pyruvate dehydrogenase and pyruvate formate lyase, enzymes that facilitate the transition to T.C.A. or fermentation by converting pyruvate to acetyl-CoA, are essential for *V. cholerae* pathogenesis in an infant mouse model^37^, and citrate metabolism intersects this metabolic node. The citrate metabolic axis in particular is a defining aspect of *V. cholerae*, commonly used to differentiate *V. cholerae* from other bacteria^38^. Citrate metabolism genes are highly conserved in *V. cholerae*^*39*^, and citrate fermentation promotes *V. cholerae* pathogenesis in an infant mouse model^26^.

Notably, *V. cholerae* citrate fermentation, along with lactate and acetate metabolism, produce carbonates as a biproduct^22^. *V. cholerae* C6706 also encodes at least three putative carbonic anhydrases (VC0586, VCA0274, and VC0058) that potentially contribute to the accumulation of environmental carbonates^40,41^. Carbonates incorporate free H+ ions to produce CO_2_, which raises pH. Additionally, OadA-1 is a decarboxylase that consumes a proton in the conversion of oxaloacetate to pyruvate, which could also contribute to environmental alkalization^28^. Both *V. cholerae* and the human host produce carbonates, and carbonate serves as an important signal in *V. cholerae* pathogenesis. Bicarbonate is a critical component of A.K.I. conditions to induce cholera toxin production *in vitro*^*42*^. It has also had more recently been shown that bicarbonate regulates the expression of the virulence regulator *toxT* and the levels of the second messenger cyclic di-GMP^43–45^. *V. cholerae* has a high tolerance for carbonates, and carbonates also inhibit the growth of *E. coli* at alkaline pH^33^. We demonstrate here that high pH and carbonates have a synergistic effect at inhibiting *E. coli* growth (Fig. 6). We postulate that mutations in the citrate fermentation pathway reduce carbonate production enough to abrogate enteric growth inhibition.

Chakrabarty et. al. first observed that *V. cholerae* inhibits growth of *E. coli* when grown on citrate containing media^21^, but we have demonstrated that external citrate is not required for this behavior (Fig. 1), similar to previous findings^22^. It is surprising that the *V. cholerae* C6706 Δ*citE* and Δ*citF* mutants were able to grow on Simmons’ citrate agar (Table 1). Despite its widespread prevalence in *Vibrio* spp., little is known about *V. cholerae* citrate metabolism^26^. In these mutant strains, citrate could be converted to oxaloacetate via the glyoxylate shunt, feeding into gluconeogenesis for the production of biomass. This pathway is mediated by isocitrate lyase (ICL), which *V. cholerae* C6706 encodes (VC0736). Consistent with this alternate citrate metabolic pathway, we simulated *V. cholerae* metabolism using a previously constructed genome-scale metabolic model based on the V52 strain^46^ with the software Optflux^47^, and found that disruption of ACLY did not impact *V. cholerae* growth *in silico* when citrate was the sole carbon source, both producing equal biomass values (0.56). This process was dependent on the glyoxylate shunt, as disruption of both ACLY and ICL resulted in no biomass production. Conversely, several of the *V. cholerae* strains examined in this study were not capable of citrate fermentation on Simmons agar. Currently, there are not genomic sequences for many of these *Vibrio* strains, but at least one strain (MZO-2) encodes citrate lyase genes with >99% protein identity to the N16961 strain, and yet does not grow on Simmons agar, suggesting that there may be differential expression of citrate metabolism genes amongst these strains. There are also examples of *V. cholerae* acquiring mutations in conserved metabolic pathways ^48^.

Citrate metabolism is not necessarily indicative that a *V. cholerae* strain inhibits enteric growth, suggesting other systems work in conjunction with citrate metabolism to create a hostile growth environment for *E. coli* and *S. flexneri* (Table 1). While we demonstrate that the citrate metabolic axis is required for *V. cholerae* C6706 inhibition of enteric growth, it is likely that other secreted molecules also contribute to this process, as we did not observe a consistent correlation amongst *V. cholerae* strains between growth on citrate, the pH of L.B. media after 48 hours, and enteric growth inhibition (Table 1); this suggests that there are multiple ways in which different *Vibrio* strains can inhibit enteric growth. There is diversity amongst *Vibrio* strains in their metabolic pathways; for example, some *V. cholerae* El Tor biotypes have developed neutral fermentation pathways, resulting in 2,3-Butanediol production, to avoid creating potentially harmful organic acids^18,19^. It is possible that certain *V. cholerae* strains produce other secondary metabolites that also contribute to enteric growth inhibition, independent of citrate fermentation. One possibility is the production of polyamines, such as cadaverine which some *V. cholerae* species produce in high abundance^49,50^, the synthesis of which consumes protons and protects *V. cholerae* from acid stress, and also inhibit enteric bacteria at high pH^51^. Further studies will reveal how other secondary metabolites contribute to *V. cholerae* inhibition of enteric bacterial growth.

This report describes an example of how *V. cholerae* C6706 utilizes a non-standard metabolic pathway to outcompete other bacteria. Specifically, we propose that creating a highly alkaline environment is a mechanism for *V. cholerae* C6706 to generate a niche; the concept of *V. cholerae* using secreted metabolites to protect its niche is not entirely novel, as a previous study has shown that *V. cholerae* releases ammonium when grown on chitin to inhibit protist grazing^52^. Creating a niche diminished of other species of bacteria would allow *V. cholerae* to suppress competing commensal bacteria and control its nutrient pool. In combination with other known systems such as the T6SS, this mechanism of competition likely contributes to the survivability, adaptability, and success of *V. cholerae* as a pathogen and prominent aquatic bacterium.

## Materials and Methods

### Strains and culture conditions

Strains included in this study are *E. coli* MG1655 and *S. flexneri* 2457T (CFS100^53^), as well as *V. cholerae* strains listed in supplementary table 1. *V. cholerae* and *E. coli* strains were grown in Lysogeny broth (L.B.) broth (1% tryptone, 0.5% yeast extract, 0.5% NaCl) at 37°C with shaking. As necessary, bacteria were grown in the presence of 50 µg/mL kanamycin, 100 µg/mL streptomycin, or 50 µ g/mL rifampicin. Cross streaking and conditioned media assays were performed using L.B. or citrate media broth (C.B.). Briefly, nutrient Broth No. 2 (Oxoid) was supplemented with NH_4_Cl (0.03%), K_2_HPO_4_ (0.5%) and sodium citrate (0.5%) and includes EGTA (3.0 mg/mL).

To determine whether a bacterium could utilize citrate as a sole carbon source, strains were grown on Simmons’ citrate agar. Ammonium dihydrogen phosphate (0.2 g/L), disodium ammonium phosphate (0.8 g/L) magnesium sulfate heptahydrate (0.2 g/L), sodium chloride (5.0 g/L) salts were added to a solution of 2.0 g/L trisodium citrate, with 1.5% w/v agar.

### Cross-streaking assay

The cross-streaking assay was performed as previously described (Bauernfeind and Burrows, 1978). Briefly, liquid from an overnight culture from the producing bacteria was streaked down the center of an agar plate (L.B. or C.B.) using a sterile cotton-tipped stick, resulting in a ∼¾” wide stripe and left to grow for 48 hours at 37°C followed by 6 hours at 4°C. The resulting growth was manually removed by scraping with a sterile cotton-tipped stick, and the remaining bacteria were killed by exposure to chloroform vapor for 30 minutes. Residual chloroform was allowed to evaporate from the plate for 30 minutes in the fume hood. Afterward, the indicator strain (for example, *E. coli* or *S. flexneri* CFS100) was streaked in a line perpendicular to the producing bacteria from an overnight culture in the same manner. The plate was then incubated for 18 hours at 37°C. For visualizing pH changes in the agar plate, an L.B. agar plate with 0.0032% w/v thymol blue was inoculated with a single streak with *V. cholerae* C6706 from an overnight culture and incubated statically for 24 hours at 37°C.

### Conditioned media assay

The concentration of our LB medium prior to bacterial growth was approximately 7.0. Both *V. cholerae* and *E. coli* or *S. flexneri* CFS100 were grown for 48 hours at 37°C and the resulting culture was centrifuged, then the supernatant was retained, and filter sterilized using 0.22 µm Millipore PVDF filters. *V. cholerae* supernatant (either diluted with saline or not) or media at specific pH was mixed 1:1 with a 1:1000 dilution of an *S. flexneri* CFS100 overnight culture in L.B. to a total volume of 100 µL in each well of a 96-well plate. *V. cholerae* supernatant produced in L.B. was used to perform the inhibition assays, using supernatant dilutions ranging from 50% to 0% (final concentration). Plates were then incubated at 37°C, shaking at 200 rpm. OD_595_ was measured at the 6-hour time point using an Opsys MR plate reader.

L.B. was adjusted to various pHs with NaOH and autoclaved. Supernatant from *V. cholerae* grown in neutral L.B. was adjusted to multiple pHs with 1 M NaOH or 1 M HCl, as appropriate, and filter sterilized using an 0.22 µm P.E.S. membrane filter. *V. cholerae* supernatant and L.B. at close pHs were combined 1:1, and their pH re-determined. The resulting media (either L.B. or 50% *V. cholerae* supernatant) were then used to perform growth inhibition assays as described above. Media were mixed 20:1 with a 1:100 inoculum of a *S. flexneri* CFS100 overnight culture. OD_595_ was measured at the 6-hour timepoint.

To determine if supernatants contained an inhibitory protein, sterile supernatants were prepared as previously described, and then these supernatants were either filtered through a 1-kDa N.M.W. ultrafiltration disc (Millipore) or treated with proteinase K. Briefly, proteinase K powder was dissolved at a concentration of 20 mg/ml in sterile 50 mM Tris (pH 8.0) and 1.5 mM calcium acetate (cite proteinase K recipe). 20 uL of this solution was added to 2 mL supernatant, and incubated at 50 °C for 20 minutes. The supernatant was then incubated at 70 °C for 10 minutes to inactivate the proteinase K, before it was used in our Conditioned media assay.

Methanol extraction of metabolites was performed as previously described, and extractions were performed in triplicate^54^. Sterile supernatants were prepared as previously described above and then 600 µL cold MeOH was added to 100 µL *V. cholerae* supernatant. The mixture was vortexed, and then centrifuged for 2 minutes at max speed at 4 °C. 100 µL mL chloroform was then added to the mixture, then vortexed and centrifuged for 2 minutes at max speed at 4 °C. 300 µL water was then added to the mixture, which was then vortexed and centrifuged for 2 minutes at max speed at 4 °C. The aqueous phase was then transferred to a new tube, and 300 µL MeOH was added. The mixture was evaporated by vacuum centrifugation, and the remaining material was resuspended in 100 µL sterile saline.

### Growth assays

For growth curves of *S. flexneri* in different concentrations of *V. cholerae* C6706 supernatant, overnight cultures of *S. flexneri* CFS100 were diluted (1:100) into 96-well plates containing L.B. with decreasing concentrations of cell-free supernatant; plates were incubated at 37°C with shaking, and the optical density at 600 nm of bacterial cultures was measured and recorded every 20 minutes for 10 hours using a BioTek Synergy H1 platereader. Data was analyzed using the R package grofit^55^, with the Richards growth model.

For growth curves of *S. flexneri* in different pH media, overnight cultures of *S. flexneri* CFS100 were diluted (1:100) into L.B. at different pH and the optical density at 600 nm of bacterial cultures was measured and recorded every hour for 7 hours. For comparing the relative growth rates of *E. coli* and *V. cholerae*, overnight cultures of *V. cholerae* or *E. coli* were diluted (1:100) into C.B. in the presence or absence of bicarbonate buffered to pH of 7 or 9. OD_600_ readings were recorded every hour. The resulting O.D.s were plotted on a graph versus time, and the portion of the graph that was linear was used to calculate a slope. The slope of *E. coli* was then divided by the slope of *V. cholerae* grown under the same growth conditions.

### In silico analysis of V. cholerae metabolism

*In silico* metabolism simulations were performed using a previously published *V, cholerae* genome-scale metabolic network reconstruction^46^ and OptFlux^47^. Although *V. cholerae* encodes a sodium/citrate symporter (VC0795), there was no reaction for citrate transport in this model^46^, so citrate exchange was modified to be cytoplasmic. We removed external boundary metabolites and used the core biomass production as our objective function and the pFBA simulation method. To simulate growth in Simmons media, default environmental conditions were used, except that the lower bound of glucose exchange was set to 0 and the lower bound of citrate exchange was set to −20. Total biomass production was limited by carbon availability under these simulated conditions.

### Statistical Analysis

All statistical analysis was performed using Graphpad prism. For supernatant inhibition assays, data were regressed using a nonlinear four parameter variable slope inhibitor model, and IC_50_ values were compared using a sum-of-squares F-test (p<0.05). For growth analysis, data were compared using ANOVA analysis (p<0.05).

## Acknowledgments

We would like to thank R.A. Finkelstein for his thoughtful contributions to this study. Work in the laboratory of S.P. is supported by the National Institutes of Health (NIH) R01 grant AI139103-01A1 and the Canadian Institutes of Health Research Operating Grants MOP-84473, MOP-137106. B.K. was a recipient of an NSERC PGS-D. Work in the laboratory of S.M.P is supported by the NIH R37 grant AI016935-35. Work in the laboratory of B.J.K is supported by the NIH R03 grant AI156432-01A1 and by startup funds and a grant from the Faculty Research and Creative Activities Award, Western Michigan University.

## Author Contributions

S.M.P., S.P., and B.J.K. contributed to conceptualization, data curation, funding acquisition, and supervision. B.K., M.E.B., J.J.B., C.C,, and B.J.K. contributed to investigation, methodology, validation, and formal analysis. B.K. and B.J.K. contributed to writing, and all authors reviewed and edited the manuscript.

## Additional Information

The authors declare no competing interests.

## Figure Legends

**Table S1**. Strains used in this study.

## References

1 Griffiths, G. L., Sigel, S. P., Payne, S. M. & Neilands, J. B. Vibriobactin, a siderophore from Vibrio cholerae. J. Biol. Chem. 259, 383–385, doi:https://doi.org/10.1016/S0021-9258(17)43671-4 (1984).

2 Hibbing, M. E., Fuqua, C., Parsek, M. R. & Peterson, S. B. Bacterial competition: surviving and thriving in the microbial jungle. Nat Rev Microbiol 8, 15–25, doi:10.1038/nrmicro2259 (2010).

3 MacIntyre, D. L., Miyata, S. T., Kitaoka, M. & Pukatzki, S. The Vibrio cholerae type VI secretion system displays antimicrobial properties. Proc. Natl. Acad. Sci. U. S. A. 107, 19520–19524, doi:10.1073/pnas.1012931107 (2010).

4 Saha, R., Saha, N., Donofrio, R. S. & Bestervelt, L. L. Microbial siderophores: a mini review. J. Basic Microbiol. 53, 303–317, doi:https://doi.org/10.1002/jobm.201100552 (2013).

5 Iwasa, Y., Nakamaru, M. & Levin, S. a. Allelopathy of bacteria in a lattice population: Competition between colicin-sensitive and colicin-producing strains. Evol. Ecol. 12, 785–802, doi:10.1023/A:1006590431483 (1998).

6 Reidl, J. & Klose, K. E. Vibrio cholerae and cholera: out of the water and into the host. FEMS Microbiol. Rev. 26, 125–139 (2002).

7 Pukatzki, S., Ma, A. T., Revel, A. T., Sturtevant, D. & Mekalanos, J. J. Type VI secretion system translocates a phage tail spike-like protein into target cells where it cross-links actin. Proc. Natl. Acad. Sci. U. S. A. 104, 15508–15513, doi:10.1073/pnas.0706532104 (2007).

8 Unterweger, D. et al. The Vibrio cholerae type VI secretion system employs diverse effector modules for intraspecific competition. Nat Commun 5, 3549–3549, doi:10.1038/ncomms4549 (2014).

9 Merrell, D. S. et al. Host-induced epidemic spread of the cholera bacterium. Nature 417, 642–645 (2002).

10 Pruzzo, C., Vezzulli, L. & Colwell, R. R. Global impact of Vibrio cholerae interactions with chitin. Environ. Microbiol. 10, 1400–1410, doi:10.1111/j.1462-2920.2007.01559.x (2008).

11 Meibom, K. L. et al. The Vibrio cholerae chitin utilization program. Proc. Natl. Acad. Sci. U. S. A. 101, 2524–2529, doi:10.1073/pnas.0308707101 (2004).

12 McCormack, W. M. et al. Evaluation of Thiosulfate-Citrate-Bile Salts-Sucrose Agar, a Selective Medium for the Isolation of Vibrio cholerae and Other Pathogenic Vibrios. J Infect Dis 129, 497–500, doi:10.1093/infdis/129.5.497 (1974).

13 Merrell, D. S., Bailey, C., Kaper, J. B. & Camilli, A. The ToxR-mediated organic acid tolerance response of Vibrio cholerae requires OmpU. J. Bacteriol. 183, 2746–2754, doi:10.1128/JB.183.9.2746-2754.2001 (2001).

14 Jonson, G., Sanchez, J. & Svennerholm, A.-M. Expression and Detection of Different Biotype-associated Cell-bound Haemagglutinins of Vibrio cholerae O1. Microbiology 135, 111–120, doi:https://doi.org/10.1099/00221287-135-1-111 (1989).

15 Nair, G. B. et al. New variants of Vibrio cholerae O1 biotype El Tor with attributes of the classical biotype from hospitalized patients with acute diarrhea in Bangladesh. J. Clin. Microbiol. 40, 3296–3299, doi:10.1128/JCM.40.9.3296-3299.2002 (2002).

16 Richardson, K., Michalski, J. & Kaper, J. B. Hemolysin production and cloning of two hemolysin determinants from classical Vibrio cholerae. Infect. Immun. 54, 415–420, doi:10.1128/iai.54.2.415-420.1986 (1986).

17 Takeya, K., Otohuji, T. & Tokiwa, H. FK phage for differentiating the classical and El T or groups of Vibrio cholerae. J. Clin. Microbiol. 14, 222–224, doi:doi:10.1128/jcm.14.2.222-224.1981 (1981).

18 Yoon, S. S. & Mekalanos, J. J. 2,3-butanediol synthesis and the emergence of the Vibrio cholerae El Tor biotype. Infect. Immun. 74, 6547–6556, doi:10.1128/IAI.00695-06 (2006).

19 Lee, D. et al. Alterations in glucose metabolism in Vibrio cholerae serogroup O1 El Tor biotype strains. Sci Rep 10, 308, doi:10.1038/s41598-019-57093-4 (2020).

20 Hu, D. et al. Origins of the current seventh cholera pandemic. Proc. Natl. Acad. Sci. U. S. A. 113, E7730–E7739, doi:doi:10.1073/pnas.1608732113 (2016).

21 Chakrabarty, A. N., Adhya, S., Basu, J. & Dastidar, S. G. Bacteriocin Typing of Vibrio cholerae. Infect. Immun. 1, 293–299, doi:10.1128/iai.1.3.293-299.1970 (1970).

22 Bhaskaran, K., Iyer, S. S., Khan, A. W. & Vora, V. C. Growth inhibition of enteric bacteria by Vibrio cholerae in nutrient media containing lactate, acetate, or citrate. Antimicrob. Agents Chemother. 6, 375–378, doi:10.1128/AAC.6.3.375 (1974).

23 Mitra, S., Balganesh, T. S., Dastidar, S. G. & Chakrabarty, A. N. Single bacteriocin typing scheme for the Vibrio group of organisms. Infect. Immun. 30, 74–77, doi:doi:10.1128/iai.30.1.74-77.1980 (1980).

24 Koser, S. A. Correlation of citrate utilization by members of the colon-aerogenes group with other differential characteristics and with habitat. J. Bacteriol. 9, 59–77, doi:doi:10.1128/jb.9.1.59-77.1924 (1924).

25 Kirchberger, P. C., Unterweger, D., Provenzano, D., Pukatzki, S. & Boucher, Y. Sequential displacement of Type VI Secretion System effector genes leads to evolution of diverse immunity gene arrays in Vibrio cholerae. Sci Rep 7, 45133, doi:10.1038/srep45133 (2017).

26 Liu, M. et al. CitAB Two-Component System-Regulated Citrate Utilization Contributes to Vibrio cholerae Competitiveness with the Gut Microbiota. Infect. Immun. 87, e00746–00718, doi:doi:10.1128/IAI.00746-18 (2019).

27 Clarke, P. H. & Meadow, P. M. Evidence for the Occurrence of Permeases for Tricarboxylic Acid Cycle Intermediates in Pseudomonas aeruginosa. Microbiology 20, 144–155, doi:https://doi.org/10.1099/00221287-20-1-144 (1959).

28 Dimroth, P., Jockel, P. & Schmid, M. Coupling mechanism of the oxaloacetate decarboxylase Na+ pump. Biochim Biophys Acta Bioenerg 1505, 1–14, doi:https://doi.org/10.1016/S0005-2728(00)00272-3 (2001).

29 Dimroth, P. & Stewart, V. Molecular Basis for Bacterial Growth on Citrate or Malonate. EcoSal Plus 1, doi:doi:10.1128/ecosalplus.3.4.6 (2004).

30 Chypre, M., Zaidi, N. & Smans, K. ATP-citrate lyase: a mini-review. Biochem. Biophys. Res. Commun. 422, 1–4 (2012).

31 Cameron, D. E., Urbach, J. M. & Mekalanos, J. J. A defined transposon mutant library and its use in identifying motility genes in Vibrio cholerae. Proc. Natl. Acad. Sci. U. S. A. 105, 8736–8741, doi:10.1073/pnas.0803281105 (2008).

32 Small, P., Blankenhorn, D., Welty, D., Zinser, E. & Slonczewski, J. L. Acid and base resistance in Escherichia coli and Shigella flexneri: role of rpoS and growth pH. J. Bacteriol. 176, 1729–1737, doi:doi:10.1128/jb.176.6.1729-1737.1994 (1994).

33 Jarvis, G. N., Fields, M. W., Adamovich, D. A., Arthurs, C. E. & Russell, J. B. The mechanism of carbonate killing of Escherichia coli. Lett. Appl. Microbiol. 33, 196–200, doi:https://doi.org/10.1046/j.1472-765x.2001.00976.x (2001).

34 Patra, T., Koley, H., Ramamurthy, T., Ghose, A. C. & Nandy, R. K. The Entner-Doudoroff Pathway Is Obligatory for Gluconate Utilization and Contributes to the Pathogenicity of Vibrio cholerae. J. Bacteriol. 194, 3377–3385, doi:doi:10.1128/JB.06379-11 (2012).

35 Wang, J. et al. Gluconeogenic growth of Vibrio cholerae is important for competing with host gut microbiota. J. Med. Microbiol. 67, 1628–1637, doi:https://doi.org/10.1099/jmm.0.000828 (2018).

36 Van Alst, A. J., Demey, L. M. & DiRita, V. J. Vibrio cholerae requires oxidative respiration through the bd-I and cbb3 oxidases for intestinal proliferation. PLoS Pathog. 18, e1010102, doi:10.1371/journal.ppat.1010102 (2022).

37 Alst, A. J. V., DiRita, V. J. & Groisman, E. A. Aerobic Metabolism in Vibrio cholerae Is Required for Population Expansion during Infection. MBio 11, e01989–01920, doi:doi:10.1128/mBio.01989-20 (2020).

38 Kobayashi, T., Enomoto, S., Sakazaki, R. & Kuwahara, S. A New Selective Isolation Medium for the Vibrio Group; on a Modified Nakanishi’s medium-TCBS agar. Nippon Saikingaku Zasshi 18, 387–392, doi:10.3412/jsb.18.387 (1963).

39 Regmi, A. & Boyd, E. F. Carbohydrate metabolic systems present on genomic islands are lost and gained in Vibrio parahaemolyticus. BMC Microbiol. 19, 112, doi:10.1186/s12866-019-1487-6 (2019).

40 Del Prete, S. et al. Biochemical properties of a new α-carbonic anhydrase from the human pathogenic bacterium, Vibrio cholerae. J. Enzyme Inhib. Med. Chem 29, 23–27, doi:10.3109/14756366.2012.747197 (2014).

41 Angeli, A., Del Prete, S., Donald, W. A., Capasso, C. & Supuran, C. T. The γ-carbonic anhydrase from the pathogenic bacterium Vibrio cholerae is potently activated by amines and amino acids. Bioorg. Chem. 77, 1–5, doi:https://doi.org/10.1016/j.bioorg.2018.01.003 (2018).

42 Iwanaga, M. & Yamamoto, K. New medium for the production of cholera toxin by Vibrio cholerae 01 biotype El Tor. J. Clin. Microbiol. 22, 405–408 (1985).

43 Abuaita, B. H. & Withey, J. H. Bicarbonate induces Vibrio cholerae virulence gene expression by enhancing ToxT activity. Infect. Immun. 77, 4111–4120, doi:10.1128/iai.00409-09 (2009).

44 Thomson, J. J. & Withey, J. H. Bicarbonate Increases Binding Affinity of Vibrio cholerae ToxT to Virulence Gene Promoters. J. Bacteriol. 196, 3872–3880, doi:doi:10.1128/JB.01824-14 (2014).

45 Koestler, B. J. & Waters, C. M. Bile Acids and Bicarbonate Inversely Regulate Intracellular Cyclic di-GMP in Vibrio cholerae. Infect. Immun. 82, 3002–3014, doi:10.1128/IAI.01664-14 (2014).

46 Abdel-Haleem, A. M. et al. Integrated Metabolic Modeling, Culturing, and Transcriptomics Explain Enhanced Virulence of Vibrio cholerae during Coinfection with Enterotoxigenic Escherichia coli. mSystems 5, e00491–00420, doi:doi:10.1128/mSystems.00491-20 (2020).

47 Rocha, I. et al. OptFlux: an open-source software platform for in silico metabolic engineering. BMC Syst. Biol. 4, 45, doi:10.1186/1752-0509-4-45 (2010).

48 Garza, D. R. et al. Genome-Wide Study of the Defective Sucrose Fermenter Strain of Vibrio cholerae from the Latin American Cholera Epidemic. PLoS ONE 7, e37283, doi:10.1371/journal.pone.0037283 (2012).

49 Merrell, D. S. & Camilli, A. Regulation of Vibrio cholerae genes required for acid tolerance by a member of the “ToxR-like” family of transcriptional regulators. J. Bacteriol. 182, 5342–5350, doi:10.1128/JB.182.19.5342-5350.2000 (2000).

50 Merrell, D. S. & Camilli, A. The cadA gene of Vibrio cholerae is induced during infection and plays a role in acid tolerance. Mol. Microbiol. 34, 836–849, doi:https://doi.org/10.1046/j.1365-2958.1999.01650.x (1999).

51 Yohannes, E., Thurber, A. E., Wilks, J. C., Tate, D. P. & Slonczewski, J. L. Polyamine stress at high pH in Escherichia coli K-12. BMC Microbiol. 5, 59–59, doi:10.1186/1471-2180-5-59 (2005).

52 Sun, S., Tay, Q. X. M., Kjelleberg, S., Rice, S. A. & McDougald, D. Quorum sensing-regulated chitin metabolism provides grazing resistance to Vibrio cholerae biofilms. ISME J 9, 1812–1820, doi:10.1038/ismej.2014.265 (2015).

53 Marman, H. E. The role of Elongation factor P in the virulence of Shigella flexneri Doctor of Philosophy thesis, The University of Texas at Austin, (2013).

54 Wessel, D. & Flügge, U. I. A method for the quantitative recovery of protein in dilute solution in the presence of detergents and lipids. Annal. Biochem. 138, 141–143, doi:https://doi.org/10.1016/0003-2697(84)90782-6 (1984).

55 Kahm, M., Hasenbrink, G., Lichtenberg-Frate, H., Ludwig, J. & Kschischo, M. grofit: Fitting Biological Growth Curves with R. J. Stat. Soft. 33, 1–21 (2010).

