## Supplementary figures and images for "*Vibrio cholerae* alkalizes its environment via citrate metabolism to inhibit enteric growth"

### Table S1

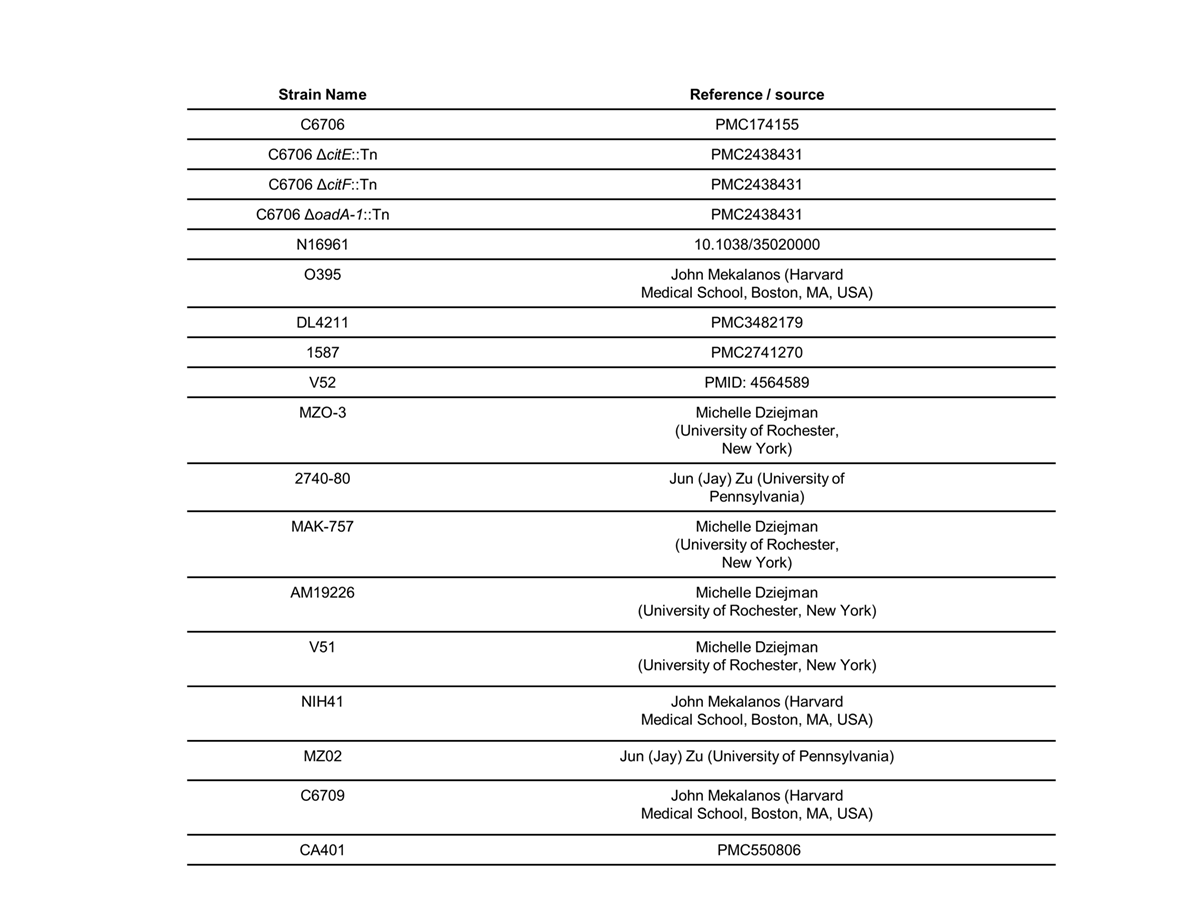
